# PSA-targeted Alpha-, Beta- and Positron Emitting Immuno-Theranostics in Murine Prostate Cancer Models and Non-Human Primates

**DOI:** 10.1101/2020.09.11.294264

**Authors:** Darren R. Veach, Claire M. Storey, Katharina Lückerath, Katharina Braun, Christian von Bodman, Urpo Lamminmäki, Teja Kalidindi, Sven-Erik Strand, Joanna Strand, Mohamed Altai, Robert Damoiseaux, Pat Zanzonico, Nadia Benabdallah, Dmitry Pankov, Howard I. Scher, Peter Scardino, Steven M. Larson, Hans Lilja, Michael R. McDevitt, Daniel L. J. Thorek, David Ulmert

**Author notes:** Correspondence to: David Ulmert, Department of Molecular and Medical Pharmacology, UCLA, Los Angeles, CA, USA. Currently an employee of GlaxoSmithKline, NJ, USA. Financial support. This study was supported in part by the Imaging and Radiation Sciences Program, US NIH grant P30 CA008748 (Memorial Sloan Kettering Cancer Center [MSKCC] Support Grant). The MSKCC Small-Animal Imaging Core Facility is supported in part by NIH grants P30 CA008748-48, S10 RR020892-01, S10 RR028889-01, and the Geoffrey Beene Cancer Research Center. We also acknowledge Mr. William H. Goodwin and Mrs. Alice Goodwin and the Commonwealth Foundation for Cancer Research, the Experimental Therapeutics Center, and the Radiochemistry & Molecular Imaging Probe Core (P50-CA086438), all of MSKCC. M.R.M.: NIH R01CA166078, R01CA55349, P30CA008748, P01CA33049, F31CA167863, the MSKCC for Molecular Imaging and Nanotechnology. D.L.J.T: NCI R01CA201035, R01CA240711 and R01CA229893. H.L., H.I.S.: NIH/NCI CCSG to MSKCC [P30 CA008748]. H.I.S.: SPORE in Prostate Cancer (P50 CA092629). H.L.: Sidney Kimmel Center for Prostate and Urologic Cancers. David H. Koch Prostate Cancer Foundation Award, Swedish Cancer Society (CAN 2017/559), Swedish Research Council (VR-MH 2016-02974), General Hospital in Malmö Foundation for Combating Cancer. D.L.J.T.: NCI R01CA201035, R01CA240711, R01CA229893. D.U., M.R.M.: DoD W81XWH-18-1-0223. D.U.: UCLA SPORE in Prostate Cancer (P50 CA092131), JCCC Cancer support grant from NIH P30 CA016042 (PI: Teitell), Knut and Alice Wallenberg Foundation, Bertha Kamprad Foundation, David H. Koch Prostate Cancer Foundation Young Investigator Award. S.M.L.: Ludwig Center for Cancer Immunotherapy (MSKCC), NCI P50-CA86438. S.-E.S.: Swedish Cancer Society, Swedish National Health Foundation, Swedish Research Council.

## Abstract

**Purpose:** Most prostate cancer (PCa) patients treated with androgen receptor (AR)-signaling inhibitors develop therapeutic resistance due to restoration of AR-functionality. Thus, there is a critical need for novel treatment approaches. Here we investigate the theranostic potential of hu5A10, a humanized monoclonal antibody specifically targeting free prostate-specific antigen (*KLK3*).

**Experimental Design:** LNCaP-AR xenografts (NSG mice) and *KLK3*_Hi-*Myc* transgenic mice were imaged with ^89^Zr- or treated with ^90^Y- or ^225^Ac-labeled hu5A10; biodistribution and subcellular localization were analyzed by gamma-counting, positron emission tomography (PET), autoradiography and microscopy. Therapeutic efficacy of [^225^Ac]hu5A10 and [^90^Y]hu5A10 in LNCaP-AR tumors was assessed by tumor volume measurements, time to nadir (TTN), time to progression (TTP), and survival. Pharmacokinetics of [^89^Zr]hu5A10 in non-human primates (NHP) were determined using PET.

**Results:** Biodistribution of radiolabeled hu5A10-constructs was comparable in different mouse models. Specific tumor uptake increased over time and correlated with PSA expression. Treatment with [^90^Y]/[^225^Ac]hu5A10 effectively reduced tumor burden and prolonged survival (p≤0.0054). Effects of [^90^Y]hu5A10 were more immediate than [^225^Ac]hu5A10 (TTN, p<0.0001) but less sustained (TTP, p<0.0001). Complete responses were observed in 7/18 [^225^Ac]hu5A10 and 1/9 mice [^90^Y]hu5A10. Pharmacokinetics of [^89^Zr]hu5A10 were consistent between NHPs and comparable to those in mice. [^89^Zr]hu5A10-PET visualized the NHP-prostate over the 2-week observation period.

**Conclusions:** We present a complete preclinical evaluation of radiolabeled hu5A10 in mouse PCa models and NHPs, and establish hu5A10 as a new theranostic agent that allows highly specific and effective downstream targeting of AR in PSA-expressing tissue. Our data support the clinical translation of radiolabeled hu5A10 for treating PCa.

## INTRODUCTION

Prostate cancer (PCa) is a leading cause of cancer death among men in the United States (1). With metastatic PCa five-year survival rates under 30%, there is a critical need for more effective systemic treatments (2); over the past decade combinations of more potent androgen receptor (AR) antagonists, androgen synthesis inhibitors and taxane-based chemotherapy were shown to improve patient outcomes and changed standards of care. None provide sustained disease control and virtually all tumors regrow, the majority of which through the restoration of AR-signaling (3).

Because of their very strong correlation with downstream AR-pathway activity, AR-governed enzymes such as human kallikrein 2 (hK2, *KLK2*) and prostate-specific antigen (PSA, *KLK3*) have been explored as both diagnostic biomarkers and therapeutic targets of PCa (4,5). These kallikrein-related peptidases are highly selectively and abundantly expressed in both healthy and malignant prostate tissues. However, common age-related pathological conditions of the prostate such as inflammation, benign age-associated enlargement and cancer result in the retrograde occurrence of PSA and hK2 in blood at levels ≈10^−6^ of those in prostate fluid. Upon their release into the extracellular compartments (e.g., blood circulation), there is little if any evidence of finding catalytically active PSA (“free” PSA, fPSA); PSA rather occurs as single or multi-chain non-catalytic, uncomplexed fPSA or has been permanently inactivated by forming a covalent complex with abundant extracellular protease inhibitors (e.g., alpha-1-antichymotrypsin, *SERPINA3*) circulating as “complexed” PSA (cPSA) (4,6,7). Measurements of the different forms of PSA and hK2 in blood can be used to predict risk of clinically significant PCa and outcome, and to monitor PCa. Recently, antibody-based methods for specific *in-vivo* targeting of fPSA and hK2 in tissue have been successfully developed and applied for *in-vivo* radio-immunotheranostics (RIT) (5,8). This approach relies on the use of high-specificity and high-affinity antibodies developed to specifically bind to the catalytic clefts of hK2 and fPSA that are uniquely exposed on the free forms of PSA and hK2, abrogates binding of the complexed form of these enzymes in the blood, and enables RIT utility in the setting of high PSA levels in the blood. This antibody technology exploits the inherent mechanism of the neonatal Fc-receptor (FcRn) to route antigen-bound monoclonal antibodies (mAbs) from recycling to lysosomal pathway compartments, resulting in internalization into target cells and accumulation of diagnostic or therapeutic radionuclides at sites of disease (9,10). Progress in antibody and small molecule design for targeted delivery and the increased availability of radionuclides with potent therapeutic properties have fueled interest in the field of targeted radiotherapy. In particular, research has focused on high linear energy transfer (LET) therapies which deliver ablative doses to cancerous cells over a small range, sparing adjacent non-targeted tissues (11,12). Radium-223 dichloride, a bone-seeking calcium mimetic for treatment of bone metastatic castrate-resistant PCa, is the first approved alpha-particle emitter (13) setting a precedent for other alpha-particle emitters undergoing clinical investigation (14). However, many questions remain, in particular the requisite properties of a RIT-construct that would enable therapeutic doses of radiation to be delivered using administration schedules that are safe and effective.

In this report, we present the results of pre-clinical studies comparing PSA-targeted RIT compounds carrying radionuclides with high or low LET. Specifically, hu5A10 - a humanized fPSA-targeting IgG_1_-mAb designed to route antigen-bound hu5A10 to the FcRn to enable its internalization into target cells - was labeled with the alpha-particle emitter Actinium-225 ([^225^Ac]hu5A10) and beta-particle emitting Yttrium-90 ([^90^Y]hu5A10), representing high- (∼100 keV/µm) and low-LET (∼0.2 keV/µm) PSA-RIT, respectively. We further evaluate the possibility of using Zirconium-89 labeled hu5A10 ([^89^Zr]hu5A10) as a companion diagnostic PET-reporter to guide therapeutic dose-planning for fPSA-RIT.

## MATERIALS AND METHODS

### Cell culture

MDAPCa2b were purchased from American Type Culture Collection. LNCaP-AR (LNCaP with overexpression of wildtype AR) was a kind gift from Dr. Charles Sawyers (15). The cell lines were cultured according to the developer’s instructions and frequently tested for mycoplasma.

### Xenograft models

All mouse studies were approved by the IACUC, MSKCC (#04-01-002). For xenograft studies, male athymic BALB/c nude mice (NU(NCr)-*Foxn1*^*nu*^; 6-8 weeks old, 20-25g; Charles River) were inoculated with LNCaP-AR or MDAPCa2b in the flank by subcutaneous (s.c.) injections of 1-5×10^6^ cells (100µl medium/100µl Matrigel). Tumors developed 3-7 weeks post-inoculation.

### Transgenic *KLK3* mouse models

Site-directed mutagenesis of APLILSR to APLRTKR at positions 4, -3, and -2 of the zymogen sequence of *KLK3* (Quick Change Lightning Mutagenesis Kit; Stratagene) enabled furin, an ubiquitously expressed protease in rodent prostate tissue, to cleave the short activation peptide at the cleavage site (−1 Arg/+1 Ile) resulting in constitutive conversion from non-catalytic zymogen to functional PSA enzyme. A transgenic mouse model was established by cloning the described construct into a SV40 T-antigen cassette downstream of the short rat probasin (pb) promoter. This construct was microinjected into fertilized mouse embryos (C57BL/6) and implanted into pseudopregnant female mice to yield the pb_*KLK3* genetically modified mouse model (GEMM). Pb_*KLK3* mice were crossed with the Hi-*MYC* model (ARR2PB-Flag-MYC-PAI transgene) to create *KLK3*_Hi-*MYC* mice, a cancer-susceptible GEMM with prostate-specific PSA expression. Integration of genes into the genome of the offspring was confirmed by Southern blot and PCR.

### Preparation of radio- and fluorescently labeled antibody constructs

Humanized 5A10 IgG_1_ mAb was developed by DiaProst AB (Lund, Sweden) and produced by Innovagen AB (Lund, Sweden) (7). The hu5A10 radio-immunoconstructs were prepared according to previously published protocols unique to the specific radionuclide (see supplementary methods). A competitive binding assay was utilized to determine the affinity of radiolabeled huA10 for fPSA; no significant loss of affinity for recombinant fPSA was noted after labeling (8).

### Biodistribution and dosimetry studies

Biodistribution studies were conducted to evaluate uptake and pharmacological distribution of fPSA targeted [^90^Y]hu5A10, [^225^Ac]hu5A10 and [^89^Zr]hu5A10. Each radioconjugate was administered intravenously and contained 30-60µg of total protein with an injected activity of 5.55MBq (150µCi; [^90^Y]hu5A10), 11kBq (300nCi; [^225^Ac]hu5A10) or 4.07MBq (110µCi; [^89^Zr]hu5A10), respectively. Biodistribution studies in LNCaP-AR tumor models were conducted at 48h, 120h and 350h for [^90^Y]hu5A10; 4h, 120h, 360h for [^225^Ac]hu5A10; and at 72h, 120h, 350h for [^89^Zr]hu5A10 post-injection (p.i.; n=3-5 per time point). To evaluate the targeting specificity of [^89^Zr]hu5A10 (350h p.i.) a blocking dose (1mg) of unlabeled hu5A10 was co-administered with the radiotracer in an additional group of LNCaP-AR mice. Biodistribution of [^89^Zr]hu5A10 was further evaluated in the MDAPCa2b model at 120h p.i. Biodistribution of all three hu5A10 radio-immunoconstructs was studied in the PSA expressing *KLK3*_Hi-*MYC* GEMM at 120h p.i. Blood was drawn by cardiac puncture immediately after mice were euthanized by CO_2_ asphyxiation. Organs/tissues were collected, rinsed briefly in water, dried on paper, weighed, and counted in a gamma-counter (Packard Instrument) for accumulation of respective radionuclide. Count data were corrected for background activity and decay and the tissue uptake for each sample was calculated by normalization to the total amount of activity injected (measured in units of %injected activity per gram [%IA/g]). Organ and tumor dosimetry calculations were performed using the Rodent Dose Evaluation Software (16,17). Briefly, pharmacokinetic data were input into a murine 3D imaging-based dosimetry platform built on accurate magnetic resonance imaging volumes in order to compute the intra- and inter-organ and tumor absorbed doses from alpha-particle, beta-particle and gamma-emission transport. The adult male mouse model and Medical Internal Radiation Dose Committee methodology (18,19) as implemented in OLINDA/EXM (20) were used to calculate the absorbed doses of [^225^Ac]hu5A10 and [^90^Y]hu5A10.

### Therapy studies

Average pre-therapeutic tumor sizes were 350±107mm^3^ (range, 268-471mm^3^) and animals were randomized into groups receiving fPSA-targeted RIT or no treatment. A single 11.1kBq (300nCi) or 18.5MBq (500µCi) activity of [^225^Ac]hu5A10 or [^90^Y]hu5A10, respectively, was injected into the tail vein in LNCaP-AR tumor-bearing mice (some with bilateral tumor grafts). In the low-LET study, tumor volume, time to nadir (TTN; i.e., time to smallest post-RIT tumor volume) and time to progression (TTP; i.e., time to two consecutive increases in tumor volume post nadir), and survival following treatment with [^90^Y]hu5A10 (n=6 mice with a total of 9 tumors) or no treatment (n=6 mice, 6 tumors) were quantified. The same parameters were recorded in high-LET [^225^Ac]hu5A10 treated LNCaP-AR xenografts (n=14 mice, 18 tumor) compared to untreated controls (n=9 mice, 16 tumors). Due to logistic issues, high-LET and low-LET treatment studies were not done in parallel. Tumor volume (*V*) was calculated by measuring the length (*L*) and width (*W*) of tumors by caliper and using the formula for a rotated ellipsoid [*V* = (*W* × 2*L*)/2]. Endpoint was defined as weight loss of 20% or a tumor diameter exceeding 15 mm. Tumor volumes were measured twice per week and survival monitored until all mice either had succumbed to tumor burden (either spontaneously or meeting criteria for euthanasia) or remained free of a visible/palpable tumor mass at the time the last mouse without complete response succumbed.

### Tissue histology and autoradiography of GEMM prostate

Bulk tissue comprised of non-separated prostate lobes, seminal vesicle and prostatic urethra was harvested following euthanasia. Tissue was embedded in optimal cutting temperature compound (Sakura) and incubated on ice for 2h before snap-freezing on dry ice in a cryomold. Sets of contiguous 5μm (autoradiography) or 100μm (fluorescence microscopy) thick tissue sections were cut with a CM1950 cryostat microtome (Leica) and arrayed onto SuperfrostPlus glass microscope slides. Autoradiographs were acquired approximately 168h after injection of *KLK3*_Hi-*MYC* GEMM with [^89^Zr]hu5A10 or [^90^Y]hu5A10 by immediately placing sectioned tissue in a film cassette against a Fuji film BAS-MS2325 imaging plate covered by 20µm thick polyvinylchloride film. The slides were exposed for 48h. Exposed phosphor plates were read by a Fujifilm BAS-1800II bio-imaging analyzer generating digital images with isotropic 50μm/pixel dimensions. For microscopy, mice were injected with 25µg Cy5.5-hu5A10 or Cy5.5-IgG1 120h before sacrifice (9). Sections stained for actin and DNA were incubated with 200μL of 10U/mL rhodamine-phalloidin (Life Sciences) in PBS for 2-3h at room temperature in a covered container and washed with PBS twice. DNA/nuclei staining was performed by incubating slides for 10 minutes in 5μg/mL DAPI in PBS, followed by a wash with PBS. Slides were air-dried and a drop of Mowiol A-48 (Calbiochem) was placed on the slide before adding a mounting cover glass. Micrographs were acquired using an Eclipse Ti-E fluorescence microscope (Nikon) equipped with a motorized stage (Prior Scientific Instruments), X-Cite light source (EXFO), and filter sets (Chroma). Images were acquired and processed using NIS-Elements AR, version 4.0 (Nikon), FIJI (National Institutes of Health), or a TCS SP8 (Leica) confocal laser scanning microscopy (Molecular Cytology Core Facility, MSKCC) (9), and MosaicJ (Phillipe Thévenaz, Biomedical Imaging Group, Swiss Federal Institute of Technology Lausanne). All fluorescent images were captured with a fixed fluorophore-dependent exposure time.

### Positron emission tomography/computed tomography (PET/CT) imaging

PET imaging experiments were conducted on a microPET Focus 120 scanner (Concorde Microsystems). Mice were anesthetized by inhalation of 1-2% isoflurane/oxygen gas mixture in order to record PET images (21). PET images were acquired at 120h after injection. The PET-bed was then moved for CT imaging using a NanoSPECT/CT (Bioscan). Data were exported in raw format, and the rigid body (three degrees of freedom) co-registration between PET and CT data was performed in Amira 5.3.3 (FEI). Details on PET and CT acquisition and analysis are provided in the supplementary methods.

### Measurement of total and free PSA

Total and fPSA were measured using a dual-label ELISA assay with 5A10 (non-humanized) as capture antibody as published previously (Prostatus PSA Free/Total PSA, Perkin-Elmer Life Sciences) (8,22, 23).

### Non-human primate (NHP) studies - Pharmacokinetics of [^89^Zr]hu5A10

The study using cynomolgus macaques was approved by the IACUC of MSKCC (#14-03-006). Animals were maintained in accordance with the USDA Animal Welfare Act and Regulations and the Guide for the Care and Use of Laboratory Animals (24,25). The animal care and use program at MSK is USDA-registered, maintains Public Health Services Assurances, and is fully accredited by AAALAC International. Binding of hu5A10 to cynomolgus PSA was confirmed by time-resolved fluorescence measurements of free and total PSA (23) in ejaculate collected from two monkeys and a purchased reference sample (Charles River) obtained by electroejaculation. The pharmacokinetics and distribution of [^89^Zr]hu5A10 was evaluated using a longitudinal PET/CT study over two weeks at the Weill Cornell Medicine – Citigroup Biomedical Imaging Center. Each animal was imaged dynamically for 60 minutes after injection, and subsequently at three more timepoints in the following ranges: 48–72h, 120–144h, and 216–312h. All details pertaining NHPs are described in the supplementary methods.

### Statistical analysis

Analysis was performed using Prism 8.0 (GraphPad). All data are presented as mean ± standard deviation (SD), unless noted otherwise. Survival, TTN and TTP were calculated using Kaplan-Meyer analysis. Statistical significance was calculated using one way-ANOVA with Bonferroni correction for multiple testing and a *P-*value ≤0.05 was considered to indicate statistically significant differences.

## RESULTS

### Macro and micro biodistribution of ^90^Y, ^225^Ac and ^89^Zr labeled hu5A10

#### Murine in-vivo/ex-vivo biodistribution studies

Targeting specificity of the radio-conjugated hu5A10 was confirmed by co-injecting [^89^Zr]hu5A10 and a 20-fold molar excess of unlabeled hu5A10, which reduced accumulation of [^89^Zr]hu5A10 (mean±SD: 7.9±0.8 %IA/g blocked, 28.6±11.8 %IA/g unblocked, p*=*0.0018) in LNCaP-AR tumors (**Supplementary Figure S1A**). Tumor uptake of [^90^Y]hu5A10, [^225^Ac]hu5A10 and [^89^Zr]hu5A10 in LNCaP-AR xenografts, in contrast to uptake in healthy organs, tended to increase over time (for [^225^Ac]hu5A10 4h vs. 120h or 350h, p≤0.0115; all other p≥0.8928) (**Supplementary Figure S1B-D**). Administration of the three different fPSA targeted radio-conjugates resulted in almost identical biodistributions; choice of chelate and radionuclide had negligible impact on tumor targeting and organ kinetics of hu5A10 (**Figure 1A**). In addition, only minimal differences were noted when comparing [^90^Y]hu5A10, [^225^Ac]hu5A10 and [^89^Zr]hu5A10 in LNCaP-AR xenografts and *KLK3*_Hi-*MYC* GEMM (**Figure 1B**). In the xenograft model, [^90^Y]hu5A10 displayed slightly longer blood retention (p=0.2634), and significantly higher bone uptake (p=0.0391) was noted for [^89^Zr]hu5A10.

**Figure 1.**
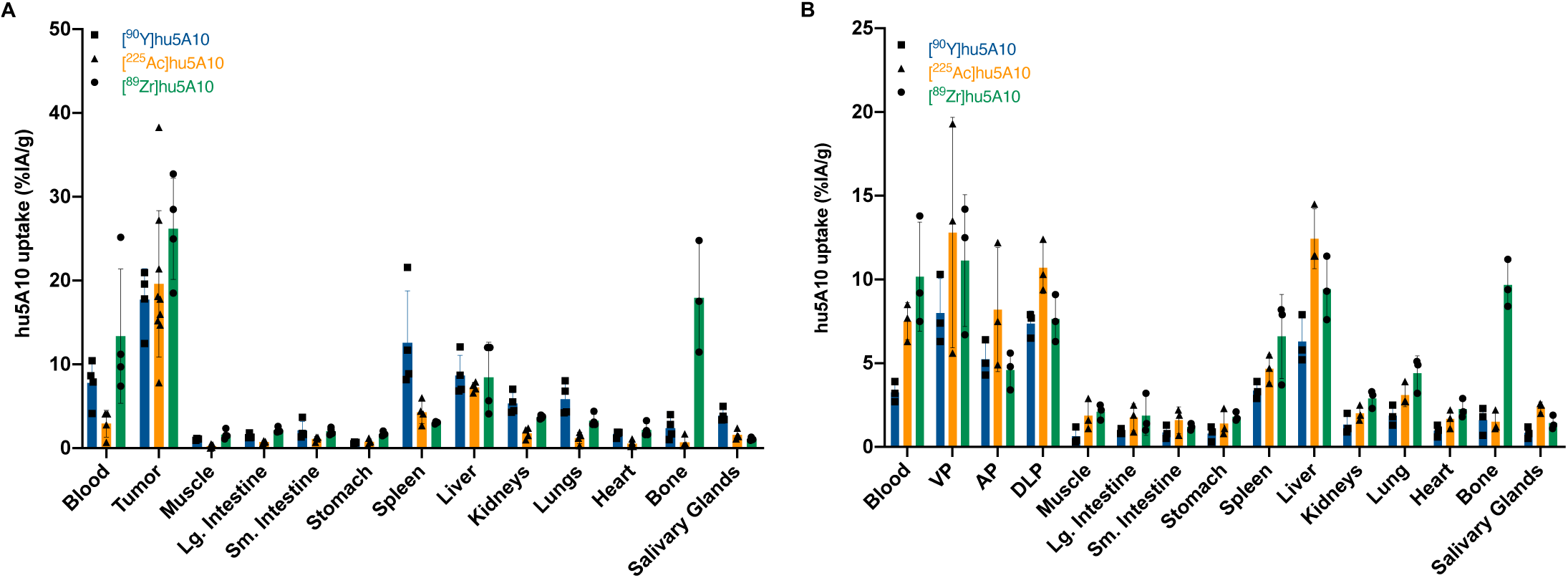
Radiolabeled hu5A10 localizes to tumor cells in PCa xenograft models. (**A**) *Ex-vivo* biodistribution data of [^89^Zr]hu5A10 (green), [^90^Y]hu5A10 (blue) and [^225^Ac]hu5A10 (orange) 120h p.i. in mice with LNCaP-AR xenograft, presented as %IA/g (n=4/group, except [^225^Ac]hu5A10, n=9 tumors). Higher retention of [^89^Zr]hu5A10 was observed in bone and blood compared to [^90^Y]hu5A10 (bone, p<0.0001; blood, p=0.0700) and [^225^Ac]hu5A10 (bone and blood, p<0.0001). Tumor uptake was higher than healthy organ uptake among all radioimmunoconjugates (p<0.0001 for all, except [^90^Y]hu5A10 tumor vs. spleen, p=0.0499, and [^89^Zr]hu5A10 tumor vs. bone, p=0.0386). (**B**) *Ex-vivo* biodistribution at 120h p.i. in *KLK3*_Hi-MYC transgenic mice (n=3/group). [^89^Zr]hu5A10 uptake in the majority of organs was comparable to [^90^Y]hu5A10 and [^225^Ac]hu5A10 (all p≥0.3823, except blood [^89^Zr]hu5A10 vs. [^90^Y]hu5A10, p=0.0001). Accumulation of all radioimmunoconjugates tended to be higher in GEMM mouse ventral prostate lobes than in anterior and dorsolateral lobes, mirroring *KLK3* expression levels (p=not significant). Columns represent mean and error bars SD. Statistical significance was calculated using one way-ANOVA with Bonferroni correction. AP-anterior prostate, DLP – dorsolateral prostate, Lg. Intestine – large intestine, Sm. Intestine – small intestine, VP – ventral prostate

hu5A10 biodistribution correlated with *KLK3* expression levels; high *KLK3*-expressing MDAPCa2b tumors had a mean [^89^Zr]hu5A10 uptake of 49.8±29.4 %IA/g compared to 26.2±6.0 %IA/g in the LNCaP-AR xenograft (p=0.0490) (**Supplementary Figure S1E**) (26). Additionally, uptake of [^90^Y]hu5A10, [^225^Ac]hu5A10 and [^89^Zr]hu5A10 in the in *KLK3*_Hi-*MYC* GEMM tended to be higher in ventral prostate lobes compared to the lower *KLK3*-expressing dorsolateral and anterior lobes (ventral vs. anterior lobes p≤0.0047 for [^225^Ac]hu5A10, [^89^Zr]hu5A10; p≥0.4546 for remainder; **Figure 1B**).

To further confirm that [^89^Zr]hu5A10 can be used as an effective theranostic surrogate for [^90^Y]hu5A10 and [^225^Ac]hu5A10, mice with LNCaP-AR tumors and *KLK3*-Hi-*Myc* GEMM were imaged with [^89^Zr]hu5A10-PET/CT. The imaging data demonstrated the capacity of [^89^Zr]hu5A10 to quantitatively visualize hu5A10 biodistribution (**Figure 2**) (8).

**Figure 2.**
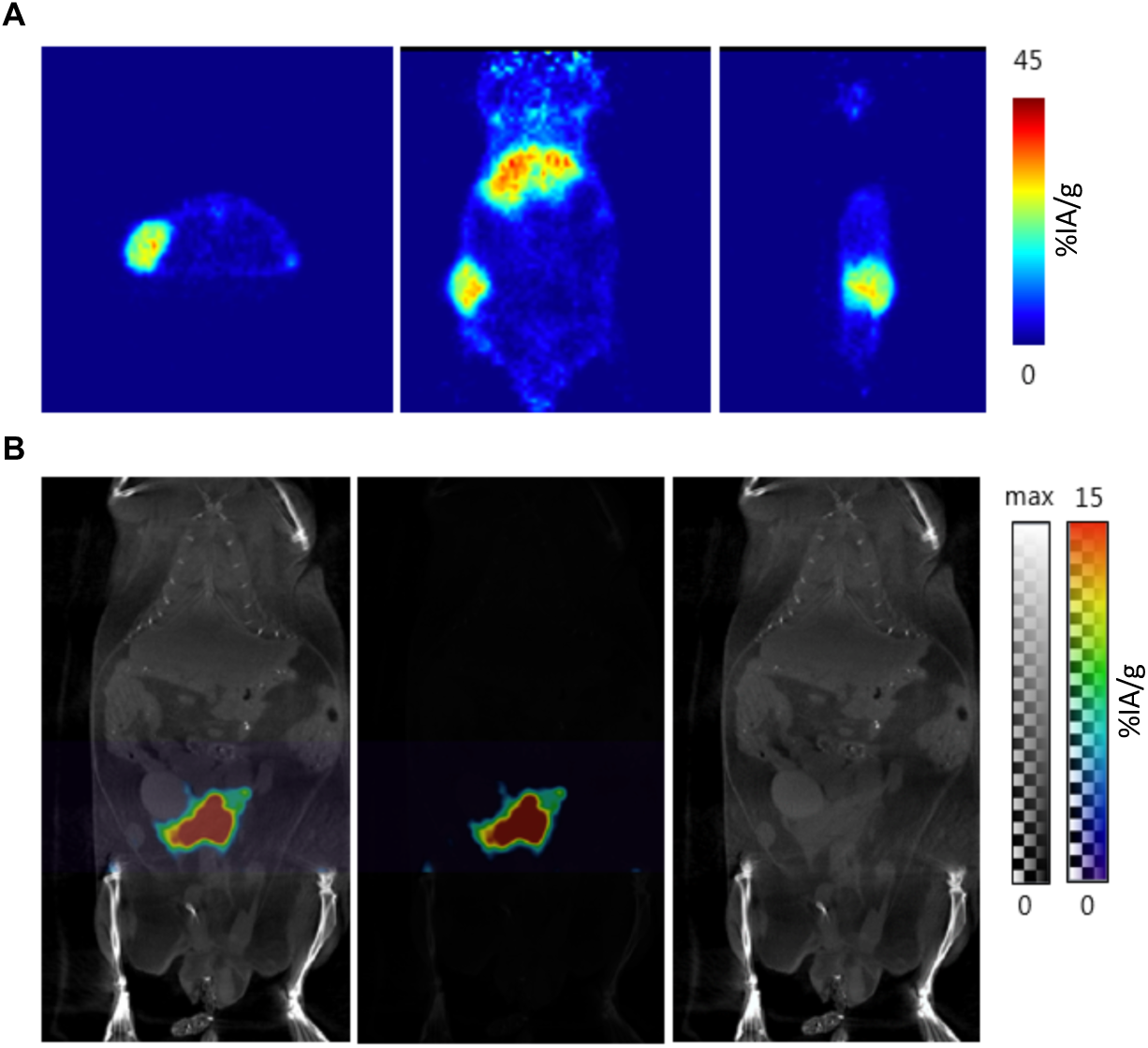
Noninvasive analysis of [^89^Zr]hu5A10 localization. Positron emission tomography was used for noninvasive and quantitative imaging of the diagnostic [^89^Zr]hu5A10 RIT conjugate. Representative images at 120h p.i. of 110µCi [^89^Zr]hu5A10 in (**A**) a mouse with LNCaP-AR xenograft (1 mouse out of 5; from left to right: transaxial, coronal and sagital view) and (**B**) the fPSA-expressing *KLK3*_Hi-*MYC* GEMM (PET/CT scan of 1 mouse out of 19; from left to right: PET/CT, PET, CT image) are shown.

#### Dosimetry

Calculating absorbed doses for [^90^Y]hu5A10 and [^225^Ac]hu5A10 from the *ex-vivo* biodistribution data showed that the tumor received the highest absorbed doses for both [^90^Y]hu5A10 (13.17Gy) and [^225^Ac]hu5A10 (9.3Gy), underlining the specific localization of the radiopharmaceuticals (**Supplementary Table S1**). The liver and the spleen were the non-targeted organs with the highest dose for animals treated with [^90^Y]hu5A10 (liver 5.36Gy, spleen 3.26Gy) and ^[225^Ac]hu5A10 (liver 1.98Gy, spleen 2.77Gy). The absorbed doses for [^90^Y]hu5A10 were higher than those calculated for [^225^Ac]hu5A10 despite the similar biodistribution profiles (**Figure 1A**). This is compatible with the higher injected activity of [^90^Y]hu5A10 (18.5MBq [150µCi], vs. 11.1kBq [300nCi] [^225^Ac]hu5A10) tolerated in this model due to the big differences in LET.

#### Autoradiography and microscopy

We further verified the organ-scale uptake with autoradiographic exposures to visualize RIT agent localization. Autoradiography of whole mounted *KLK3*_Hi-*MYC* GEMM prostate sections showed correlations between regions of increased [^89^Zr]hu5A10 or [^90^Y]hu5A10 activity and lobes expressing high levels of PSA with little if any difference between the therapeutic and diagnostic radio-conjugates (**Supplementary Figure S2**). No uptake was noted in adjacent tissues lacking PSA expression, such as seminal vesicles and urethra.

As autoradiographic investigation is limited by phosphor-sheet composition resolution, we conducted high resolution sub-cellular localization studies by preparing a fluorescent (Cy5.5) immunoconjugate for comprehensive immunofluorescent investigations of prostate tissue from *KLK3*_Hi-*MYC* GEMM following systemic administration of Cy5.5-hu5A10 or isotype control. Cy5.5-hu5A10 localized to the prostate epithelium and lumen of the glandular prostate ducts (**Figure 3A**); the concentration of the fluorescently-labeled tracer at these sites was specific, as control non-specific human IgG1 antibody was not detected (**Figure 3B**). These data confirm previous reports using antibodies targeting prostate kallikreins and are consistent with prior observations showing that FcRn facilitates transcytosis of unbound huIgG1 (hu5A10) from blood circulation to a glandular lumen, while huIgG1 bound to antigen (hu5A10:fPSA) is subjected to endosomal maturation before fusing with lysosomes (9,10). Underlining the internalization of the fPSA-specific antibody conjugate (8), diffuse signal was also present in the seminal vesicles themselves. To ascertain luminal cell transcytosis of Cy5.5-hu5A10 we used similarly processed thicker tissue sections and imaged by confocal microscopy (**Figure 3C, D**). Internalization of the Cy5.5-hu5A10 was observed as foci in luminal cells throughout the sub-region of the dorsolateral prostate.

**Figure 3.**
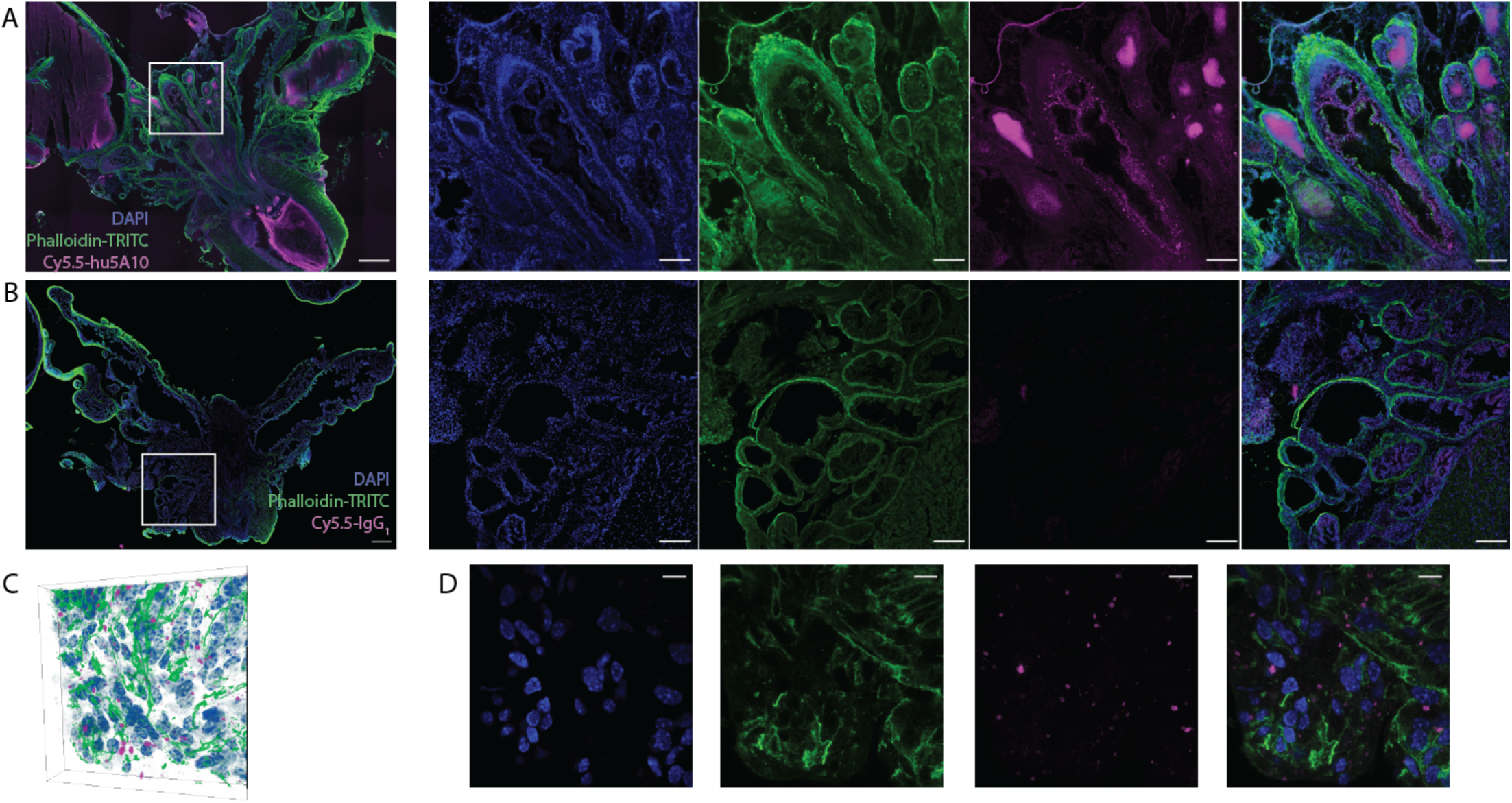
Subcellular localization of hu5A10. (**A**,**B**) Representative tissue section (100µm) through the prostate gland and seminal vesicles of a *KLK3*_Hi-*MYC* mouse at 120 h p.i. of either PSA-targeted antibody (Cy5.5-hu5A10; 1 out of 5 is shown) (**A**) or control (Cy5.5-IgG1; 1 out of 5 is shown) (**B**); composite image followed by individual channels from right to left. Concentration of the PSA-targeting Cy5.5-hu5A10 is noted in luminal structures as well as diffusely in the seminal vesicles. No signal in the Cy5.5 channel is noted following Cy5.5-IgG1 administration. Scale bar for composites is 500µm; 200µm for panels. (**C**,**D**) Microscopy three-dimensional data (**C**) of the distribution of the fluorescent Cy5.5-hu5A10 throughout the dorsolateral prostate (1 out of 5 is shown). The center slice through the prostate volume in C is shown for each channel (antibody, cell nuclei and actin) (**D**). Transport of Cy5.5-hu5A10 through the luminal PSA-expressing cells is noted, indicating internalization. The scale bar represents 40µm.

#### Treatment efficacy of low- and high-LET fPSA-targeted RIT

The efficacy of fPSA-targeted RIT with [^90^Y]hu5A10 (low-LET) and [^225^Ac]hu5A10 (high-LET) was examined in LNCaP-AR xenografts. Compared to the non-treated animals, both [^90^Y]hu5A10 and [^225^Ac]hu5A10 significantly increased disease control. The effects of [^90^Y]hu5A10 on tumor volume seemed to be more immediate than [^225^Ac]hu5A10 (**Figure 4A**) with a median TTN of 7 days for [^90^Y]hu5A10 and of 38 days for [^225^Ac]hu5A10 treated mice (p<0.0001; **Figure 4B**). However, the treatment effects of [^90^Y]hu5A10 were less sustained than [^225^Ac]hu5A10 resulting in median TTP of 24 days ([^90^Y]hu5A10) versus 72.5 days ([^225^Ac]hu5A10; p<0.0001) (**Figure 4C**). This difference entailed a median survival of 188 days with [^225^Ac]hu5A10, which was significantly longer than the survival of [^90^Y]hu5A10 treated animals (64 days, p*=*0.0009) (**Figure 4D**). At the end of the study, 7/18 animals (38.9%) receiving the high-LET treatment exhibited complete responses with unpalpable tumor burden, whereas only 1/9 animals (11.1%) in the low-LET group presented with a similar outcome. However, both treatments conferred an overall survival benefit when compared to untreated controls with a median survival of 32 days (p≤0.0054).

**Figure 4.**
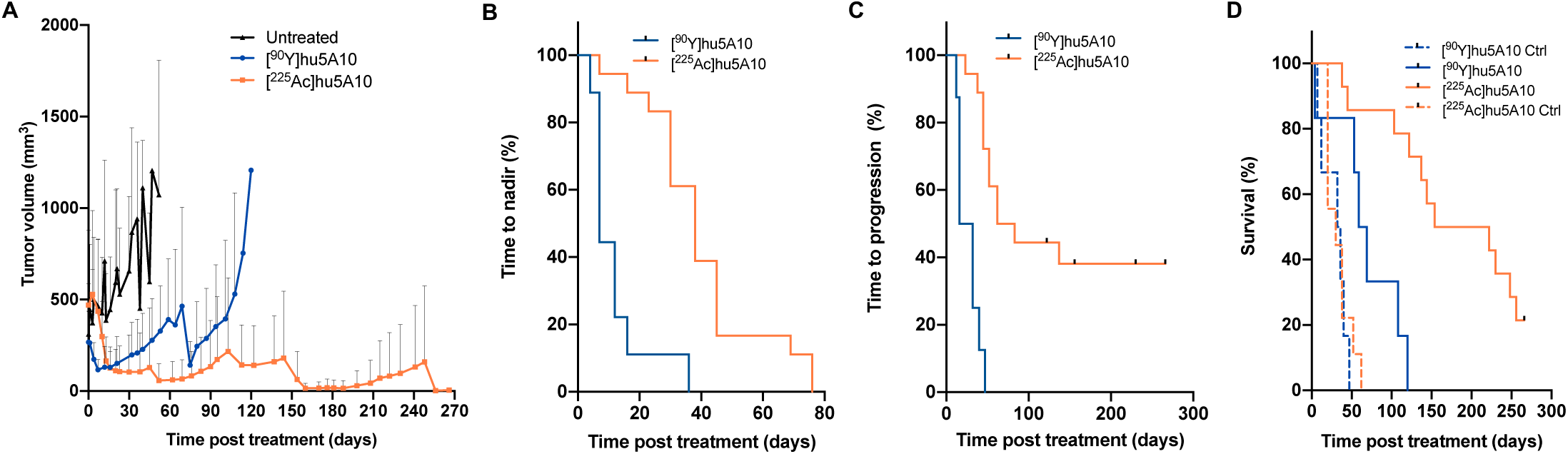
Low- and high-LET radioimmunotherapies have different outcomes in survival and tumor volume. Mice with LNCaP-AR s.c. tumors were treated with a single activity of 300nCi (11.1kBq) [^225^Ac]hu5A10, 18.5MBq (500µCi) [^90^Y]hu5A10, or left untreated. (**A**) Tumor volumes. Data represent mean + SD of n=22 tumors (control), n=9 tumors ([^90^Y]hu5A10), and n=18 tumors ([^225^Ac]hu5A10). (**B**) TTN. Treatment with [^90^Y]hu5A10 (n=9 tumors) demonstrated overall more rapid effects on tumor volume than [^225^Ac]hu5A10 (n=18 tumors), with median TTN of 7 days and 38 days, respectively (p<0.0001). (**C**) TTP was 24 days following treatment with [^90^Y]hu5A10 (n=9 tumors) and 72.5 days following [^225^Ac]hu5A10 (n=18 tumors; p<0.0001). (**D**) Survival following [^90^Y]hu5A10 (blue line; n=6 mice), [^225^Ac]hu5A10 (orange line; n=14 mice), or no treatment (dashed lines; n=6-9 mice). Alpha-RIT increased survival from 32 days (control) to 188 days (p<0.0001), while beta-RIT increased survival to 64 days (p=0.0054 vs control; p=0.0009 vs. [^225^Ac]hu5A10).

#### In-vivo biodistribution and imaging studies in NHP

Binding to PSA in seminal plasma confirmed that hu5A10 can detect cynomolgus monkey PSA. Measured PSA levels in a reference sample of seminal fluid were 44.0±8.66 ng/mL, while 4.25±1.84 ng/mL and 28.8±9.62 ng/mL fPSA were detected in samples from monkeys included in this study; the latter samples were of lower quality and suspected to having been subject to degradation *in-vitro*. Despite these very low levels relative to man, we endeavored to investigate whether hu5A10-uptake could be visualized in a non-human primate model. [^89^Zr]hu5A10 (0.2mg/kg; 115±11MBq) was administered to three adult, male cynomolgus macaques and imaged longitudinally by PET/CT over two weeks. No adverse events or physiologic responses were observed. All observed parameters (e.g., blood pressure, heart rate, body temperature) remained in the normal range and unchanged during injection and throughout the imaging procedure. All NHP subjects recovered normally from anesthesia on the day of the initial administration of [^89^Zr]hu5A10 and on subsequent imaging days. The pharmacokinetic behavior of [^89^Zr]hu5A10 was similar between the three NHP subjects studied (**Figure 5A-G, Supplementary Figure S3**) and the aggregate data were generally very consistent with the slow bulk tissue clearance observed in mice; in the absence of a tumor sink, the liver had the highest relative uptake in both species. [^89^Zr]hu5A10 rapidly entered the blood pool and distributed into most tissues within the first hour followed by a slow clearance over several weeks from most tissues. The NHP prostate gland was visualized clearly by [^89^Zr]hu5A10 PET/CT over the entire observation period with SUV_*mean*_ reaching 5.9±2.3 at 1h p.i. and decreasing thereafter to SUV_*mean*_ 4.5±1.4 (2-3 days), 2.9±1.2 (5-6 days) and 2.0±0.6 (9-13 days). The time-activity-curves (**Figure 5E**) demonstrated significant uptake in epididymis, less so in testis and minimal uptake in the seminal vesicles. The early 1h pattern of uptake in the epididymis paralleled the blood pool and did not follow the slower uptake curve in prostate and testicles. Radioactivity transit into bladder and intestine was insignificant (SUV_*mean*_<1); the majority of the off-target radioactivity residualized in liver and was not excreted. Spleen accumulation peaked at 30 minutes (SUV_*mean*_ 7.5±0.5) and dropped to SUV_*mean*_ of 3.9±0.4. Kidney uptake reached an early plateau (SUV_*mean*_ 4.8±0.5) several minutes after injection; the spike in the time-activity-curve 30 seconds after injection was due to a significant spillover effect from the adjacent large vessels as the high initial input activity transits in close proximity (**Supplementary Figure S3**). A measure of uptake in bone was determined in four lumbar vertebrae (L1–L4) which reached a SUV_*mean*_ of 2.3±0.7 on the day of administration, but stabilized at a low plateau of 1.2–1.3 SUV_*mean*_ afterwards. There was negligible accumulation in brain, lung, or salivary glands as anticipated, so these tissues were not included in the volume-of-interest analysis.

**Figure 5.**
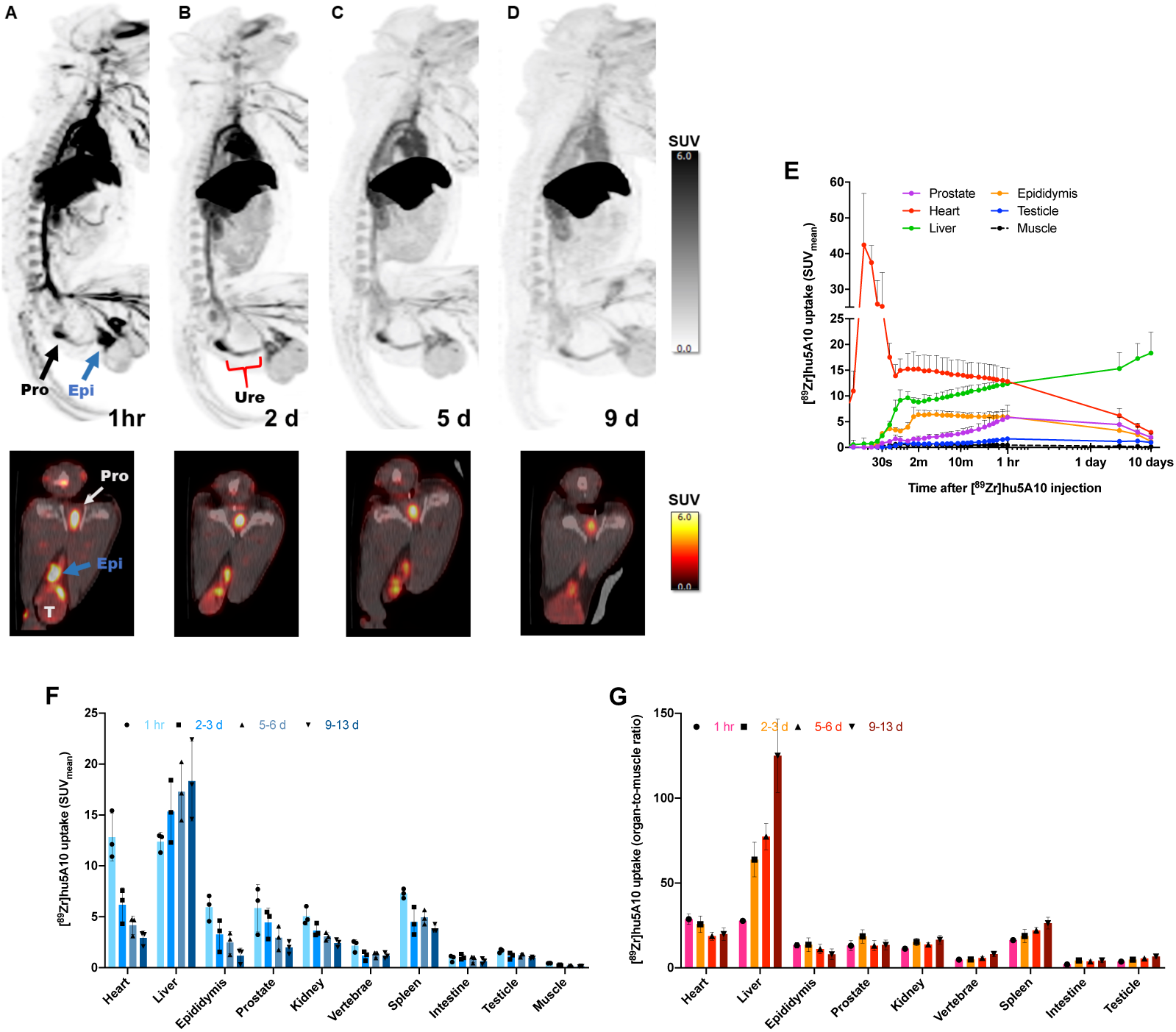
Longitudinal [^89^Zr]hu5A10 PET imaging in NHP. Male cynomolgus macaque were intravenously administered 115±11MBq (3.1±0.3mCi) of [^89^Zr]hu5A10 at 0.2mg/kg for longitudinal immune-PET imaging. (**A-D**) *Top panel*: Maximum intensity projection (MIP) whole-body images of a 14 years-old, 10.7 kg male macaque at 1 hour (sum of 0-60 minutes; (**A**), 2 days (**B**), 5 days (**C**) and 9 days; (**D**) post-injection. Image thresholds are at SUV=6. Arrows denote focal uptake in prostate (black), epididymis (blue) and testicles (T). *Bottom panel*: Transaxial PET/CT slice showing the same structures. (**E**) Time-activity-curves of [^89^Zr]hu5A10 (n=3). SUV_*mean*_ (mean+range) were calculated for each organ volume-of-interest and plotted over time. Time-activity-curves for additional organs are shown in Supplementary Figure S3. (**F**) PET-guided biodistribution of [^89^Zr]hu5A10 (n=3) at 1 hour, 2-3 days, 5-6 days, and 9-13 days. (**G**) PET derived organ-to-muscle ratio (SUV_organ_/SUV_muscle_) of [^89^Zr]hu5A10 (n=3). Columns represent mean and error bars SD.

## DISCUSSION

In this study, we present fPSA as a novel target for alpha- and beta-RIT in PCa representing high and low LET therapy, respectively and introduce a viable companion diagnostic to report pharmacokinetic accumulation and clearance. Therapy with fPSA-targeted mAb hu5A10 IgG_1_ resulted in a significant reduction in tumor burden and improvement in survival with both alpha- and beta-particle emitting radionuclides. [^89^Zr]hu5A10 is a highly specific and complementary imaging agent for diagnosis and dose planning. Yttrium-90, Actinium-225, and Zirconium-89 labeled hu5A10 localized in a *KLK3*-dependent manner in murine PCa models and primates ([^89^Zr]hu5A10) with similar pharmacokinetics and biodistribution profiles. Despite the observed similarity in radiotracer uptake, [^225^Ac]hu5A10 treatment prolonged progression-free and overall survival compared to [^90^Y]hu5A10 and offered larger overall reduction in tumor volume demonstrating important therapeutic differences between high- and low-LET therapeutic radionuclides. Additionally, hu5A10 demonstrated accumulation in target cells as a result of FcRn-mediated internalization of these RIT complexes, increasing therapeutic benefit and preventing off-target tissue damage that may accompany alpha-radionuclide therapy.

Efforts to exploit biomarkers of PCa for targeted radionuclide theranostics have led to many promising advances in management and treatment. However, congruent strategies for administering therapeutics and monitoring disease response are desperately needed in the field given the rates of PCa recurrence and acquired resistance to conventional therapies (27,28). Given the role of the AR pathway in tumor development and progression, PCa treatments have focused on targets downstream of AR activity such as prostate-specific membrane antigen (PSMA) and six-transmembrane epithelial antigen of the prostate-1 (STEAP1) with varying success. Radionuclide therapies targeting PSMA have shown some successes in disease management but are limited in their use for imaging of AR pathway activity because of the inverse correlation to AR activation, a feature that might render PSMA-targeted therapies susceptible to development of resistance upon AR reactivation and subsequent loss of PSMA expression (29). Furthermore, alpha-particle therapies targeting PSMA are often hindered by dose-limiting toxicities (see below) (30). Approaches directed to STEAP1, a membrane protein highly upregulated in multiple cancer types, seemingly demonstrated AR-dependent STEAP1 expression in the prostate (31,32); however, studies investigating STEAP1 regulation have shown both AR-dependent and independent regulation in various preclinical PCa models, suggesting a more nuanced role for AR in regulating STEAP1 expression (33,34). PSA has been primarily used as a PCa biomarker in the blood, yet serum levels of total PSA do not reliably discern PSA from malignant vs. healthy tissue nor provide reliable correlations with AR expression. Instead, targeting fPSA before it has been released and complexed in the blood is a unique approach and of significant clinical interest. The possibility to target fPSA with AR pathway-dependent localization while avoiding circulating PSA in the blood was initially reported by us using murine 5A10 (8). Subsequent modification of the Fc-region of the mAb (9) enabled the tissue-specific internalization of hu5A10 paving the way for use of hu5A10 in radioimmunotheranostics.

Internalization of a radioimmunoconjugate by the target cell is critical to the delivery of radionuclide therapies to ensure that decay of the radionuclide is on target (35). Given the short path length and high LET of alpha-particles, success of [^225^Ac]hu5A10 particularly depends upon internalization mechanisms; internalization of Actinium-225 also ensures that the radionuclide daughters remain in the cell and contribute to the therapeutic response (12). As shown by confocal microscopy, our data suggest that internalization and uptake of hu5A10 into the prostate epithelium is facilitated by IgG binding to FcRn. FcRn mediates IgG recycling as well as transport and transcytosis across epithelial cells through IgG:FcRn binding in low pH conditions allowing hu5A10 to reach target tissues and avoid washout (10). FcRn-mediated internalization of the hu5A10 complex resulted in significant accumulation in tumor cells while healthy organs had minimal uptake, thus mitigating off-target effects and sparing healthy tissue. Other studies of alpha-particle based radionuclide therapies in PCa have found dose-limiting accumulation of radionuclide in salivary glands or kidneys with unexpectedly higher uptake in healthy organs than beta-emitting RIT with the same biomarker (30,36,37,38). While initial uptake in healthy tissue was noted at 4h p.i., clearance of [^225^Ac]hu5A10 from non-target tissue occurred within 120h p.i. resulting in high tumor-to-tissue ratios. Clearance from the liver was not as pronounced in the LNCaP-AR xenograft model indicating that hepatic toxicity could be a potential limitation of [^225^Ac]hu5A10. This could be a result of the prevalent FcRn expression in the liver, a function critical for regulating albumin homeostasis (39).

The results with Actinium-225 and Yttrium-90 yield critical distinctions for PCa RIT. While beta-emitting [^90^Y]hu5A10 had a more immediate effect on tumor volume, treatment with [^225^Ac]hu5A10 sustained tumor suppression and provided a significant increase in median survival time (**Figure 4**). The faster response time seen in Yttrium-90 treatment could be attributed to the difference between the chosen radionuclides in half-life and path length. Yttrium-90 has a half-life of 2.5 days as opposed to Actinium-225 at 9.9 days and thus, could deliver the absorbed dose with a higher absorbed dose rate than Actinium-225 (38,40). In addition, the 5mm path length of Yttrium-90 (38) provided a larger immediate effect on the PCa tumors given their average size of approximately 200mm^3^ at treatment initiation. While the 50µm path length of Actinium-225 may not have an immediate effect on tumor burden, the emission of 4 alpha-particles and 2 beta-particles upon decay and ability to cause double-stranded breaks (40) likely induced irreversible cell damage and prevented PCa tumor progression. These findings are in agreement with the greater relative biological effectiveness of Actinium-225 compared to Yttrium-90 (38).

Previous work demonstrated a similar strategy to target hK2, an AR pathway-dependent peptidase closely related to PSA. Treatment with anti-hK2 [^225^Ac]hu11B6 in PCa xenografts and Hi-*Myc KLK2*-expressing GEMM demonstrated alpha-particle damage-induced upregulation of AR and hK2 expression, leading to further uptake of the hK2-targeting radioimmunoconjugate (5). Double-strand DNA breaks by alpha-emitting [^225^Ac]hu5A10 may induce the same AR activation and increase in PSA expression, leading to its intratumoral accumulation and contributing to sustained treatment effects following [^225^Ac]hu5A10 RIT compared to [^90^Y]hu5A10.

Labeling the hu5A10 construct with the PET-tracer Zirconium-89 provided a strategy for noninvasive imaging of PCa cells and complemented the PSA-targeted RIT described. Earlier work demonstrated transient localization of mouse monoclonal 5A10 in an AR-dependent manner to both prostate lesions and metastatic disease of prostatic origin (8). In this study, a transgenic *KLK3*-expressing Hi-*Myc* GEMM showed increased accumulation of [^89^Zr]hu5A10 in the ventral prostate lobes compared to adjacent dorsolateral and anterior prostate lobes, correlating with higher expression levels of PSA in this region. Additionally, MDAPCa2b xenografts showed higher [^89^Zr]hu5A10 uptake correlating to the higher *KLK3* expression in MDAPCa2b vs. LNCaP-AR demonstrating the sensitivity of hu5A10 accumulation to reflect changes in PSA expression. Using the described radioimmunoconjugate, [^89^Zr]hu5A10 PET could be used to monitor changes in AR pathway activity.

Imaging studies in primates show that [^89^Zr]hu5A10 is well tolerated at the 0.2mg/kg dose level and good image quality is obtained with 115MBq (3mCi) on a clinical PET/CT scanner. This activity was high enough to ensure that imaging up to two weeks post injection was feasible. Importantly, prostate visualization becomes clear within the first hour of imaging and the prostate is clearly discernable at all imaging timepoints over 2 weeks, which is clear from the transaxial imagery (**Figures5A-D)**. The highest quality [^89^Zr]hu5A10 prostate scan (signal-to-background) was obtained 2–3 days post injection, which is within the first half-life of Zirconium-89. This suggests that long-term imaging is likely unnecessary and injected activity might be reduced without degrading diagnostic value. Our *in-situ* assessments of PSA obtained in ejaculates confirmed previously reported findings that PSA levels in cynomolgus monkeys are approximately 5,000-fold lower than in humans (41,42). In addition, cynomolgus monkeys express two alternative splice variants of PSA and only one of these products constitutes an intact catalytic triad. It is also unclear whether the minor amino acid sequence dissimilarity in the mature cynomolgus monkey PSA results in structural differences that affects enzymatic activity or affinity of hu5A10. Previously translated PCa radioimmunotheranostics such as 7E11 and J591 have suffered from low uptake in soft tissue tumors and off-target binding in kidneys (3,43). The target-specific accumulation of [^89^Zr]hu5A10 in NHP, despite the low levels of fPSA, is highly encouraging for future clinical translation to humans. Our results also showed that hu5A10 has an extended blood circulation time, a feature shared by all IgG1 with intact FcRn binding. It is not unlikely that this will result in dose limiting myelosuppression when utilizing hu5A10 labeled with beta-emitting radionuclides, such as Lutetium-177 and Yttrium-90. However, these side effects are commonly transient and can be successfully limited by fractionated dosing strategies or utilizing radionuclides emitting alpha-particles (22,44). We also noticed uptake in epididymis, vas deferens and the seminal vesicles; measurable PSA levels in lysates of these organs have been reported in humans, albeit at very low levels in relation to prostate tissue (45). It is currently unknown whether the relative PSA expression in these organs compared to prostate are different in NHPs.

In summary, this study presents a complete preclinical evaluation in multiple rodent disease models and NHPs of fPSA-targeting hu5A10 IgG_1_. Our results show that this novel PCa theranostic is highly applicable and effective for radioimmunotherapy and diagnostic PET imaging. The study further demonstrates the different therapeutic outcomes from beta- and alpha-emitting versions of the hu5A10 mAb; while utilization of the former radionuclide results in more instant therapeutic events, the latter shows slower but more protracted anti-tumor effects with higher chance of cure.

## TRANSLATIONAL RELEVANCE

This study presents a novel theranostic approach for the treatment of prostate cancer utilizing the humanized monoclonal antibody hu5A10 to target androgen receptor-regulated free prostate-specific antigen (fPSA). Using murine models of prostate cancer, we developed and evaluated alpha and beta emitting radionuclides for radioimmunotherapy with radiolabeled hu5A10. Radioimmunotherapy targeting fPSA is a highly effective therapeutic approach for improved survival and to facilitate local disease control and complete response. We report important differences in the therapeutic outcome following alpha- vs. beta-radioimmunotherapy, with alpha-radioimmunotherapy exhibiting delayed but more potent and more sustained anti-tumor effects. Positron emission tomography with [^89^Zr]hu5A10 complements fPSA-radioimmunotherapy by enabling diagnosis and patient stratification for subsequent fPSA-targeted therapy. Notably, [^89^Zr]hu5A10 PET-uptake in the prostate of non-human primates was high despite several thousand-fold lower fPSA levels in monkeys in comparison to humans, further supporting clinical translation of hu5A10.

## Acknowledgements

We thank Muc Du, Simon Moritz, Ed Fung, Heather Martin (CBIC), and Brad Beattie, Daniel LaFontaine (MSKCC) for their excellent assistance with animal imaging/care, and data acquisition/processing, respectively.

