## Supplementary Methods for "PSA-targeted Alpha-, Beta- and Positron Emitting Immuno-Theranostics in Murine Prostate Cancer Models and Non-Human Primates"

**Radio- and fluorescent labelling of antibodies**

***General Radiochemistry.*** Reagents and buffers were tracemetal grade or equivalent, plastic pipette tips were PCR grade and microcentrifuge (reaction) tubes were acid-washed. Buffers were purified of residual metal-ion contamination in batch with 5% w/v ion exchange resin (Chelex 100, Bio-Rad) and adjusted to final pH while in contact with resin. Radiochemical reactions were monitored by radio-instant thin-layer chromatography (Radio-ITLC) with a (stationary phase of silica gel impregnated paper (Gelman Science) and quantitated on a Bioscan AR-2000 scanner under P10 (10% methane / 90% argon) gas flow or by direct gamma counting. Generally, reaction conversions were nearly quantitative, so the mass of radiometallated hu5A10 was assumed to be proportional to radioactivity for yield and specific activity calculations.

***[$^{225}$Ac]hu5A10.*** $^{225}$Ac (ORNL) was conjugated to the hu5A10 antibody using a 2-step labeling procedure (1,2). Activity was measured at secular equilibrium with a Squibb CRC-17 Radioisotope Calibrator (E.R. Squibb and Sons) set at 775 and multiplying the displayed activity value by 5. The radiochemical purity of the final product, [$^{225}$Ac]hu5A10, was determined using ITLC and two different mobile phases. Mobile phase I was 10 mM ethylenediaminetetraacetic acid and phase II was 9% sodium chloride/10 mM sodium hydroxide. The strips were counted in a gamma-counter (Packard Instrument) using a 370–510 KeV energy window. The purified radio-immunoconstruct was formulated in a solution of 1% human serum albumin (HSA; Swiss Red Cross) and 0.9% sodium chloride (Normal Saline Solution) for intravenous injection.

***[$^{89}$Zr]hu5A10.*** Radiolabeling of hu5A10 with Zirconium-89 was performed by one-step labeling of pre-conjugated DFO-hu5A10. To conjugate DFO (deferoxamine) to hu5A10, 1–2mg of antibody was buffer exchanged with reaction buffer, pH 8.5 0.5M HEPES using an Amicon Ultra-4 centrifugal 50kD MWCO filter (Millipore) at 4,000xg. Ten molar equivalents of a DMSO stock solution of p-SCN-Bn-DFO (B-705, Macrocyclics) were added in three aliquots to hu5A10 and incubated 60 minutes at 37°C. The DFO-hu5A10 conjugate was purified by four buffer exchanges in another Amicon unit with pH 7.5 0.5M HEPES buffer. Here, the final concentrate was used immediately in the radiolabeling. DFO-hu5A10 was stable at 4°C and could be used weeks later. $[^{89}Zr]Zr^{4+}(C_2O_4^{2-})_2$ in 1M oxalic acid was used as supplied (MSKCC RMIP Core or 3D Imaging LLC). $^{89}Zr$ sources were measured in a CRC-55tR dose calibrator using setting 517 (3). The $^{89}Zr$ stock was neutralized by adding small aliquots of 1M $Na_2CO_3$ until neutral by narrow range pH paper. After 5 minutes the salts, if observed, were removed by centrifugal filtration (Costar 8169, Corning). The clear $^{89}Zr$ solution was added to the DFO-hu5A10 conjugate, mixed gently by pipetting and incubated at room temperature for 1 hour. Radio-ITLC in 50mM diethylenetriamine pentaacetate (DTPA) was performed to confirm consumption of free $^{89}Zr$. DTPA (20 µL, 50mM) was added to the reaction mixture to chelate any remaining free $^{89}Zr$. Crude $^{89}Zr$-DFO-hu5A10 ($[^{89}Zr]$hu5A10) was purified on a PD-10 column (GE Healthcare) gravity eluted with 1% HSA in 0.9% saline and evaluated by radio-ITLC for radiochemical purity. Radiolabeling was efficient at a range of specific activities and generated high purity radioimmunoconjugate: scale, 1.96±0.78 mg (range, 1.20-3.20 mg); yield: 176.80±27.89 MBq (range, 141-218 MBq, 3.81-5.89 mCi) with 91.42±3.13 %conversion (range, 88.00-95.10 %conversion); radiochemical purity: 99.22±0.68 % (range, 98.30-99.80 %); specific activity of 137.20±39.42 MBq/mg (range, 96–195 MBq/mg, 2.6–5.3 mCi/mg) (mean±SD, n= 5).

***$[^{90}Y]$hu5A10.*** Radiolabeling of hu5A10 with Yttrium-90 was performed by one-step labeling of pre-conjugated DOTA-hu5A10. To conjugate DOTA to hu5A10, 1 mg of antibody was buffer exchanged with reaction buffer, pH 8.7 0.1M $NaHCO_3$ using an Amicon Ultra-4 centrifugal 50kD MWCO filter (Millipore). Ten molar equivalents of a DMSO stock solution of p-SCN-Bn-DOTA (B-205, Macrocyclics) was added in three aliquots to the hu5A10 and incubated 35 minutes at 37°C. The DOTAhu5A10 conjugate was purified by four buffer exchanges in another Amicon unit with pH 7.2 0.5M $NH_4OAc$ buffer. Here, the final concentrate was used immediately in the radiolabeling. DOTA-hu5A10 was stable at 4°C and could be used weeks later. $[^{90}Y]Cl_3$ in 0.05M HCl was used as supplied from PerkinElmer. Yttrium-90 sources were measured in a CRC-55tR dose calibrator using setting 48x10 (4) and positioned in the bottom center of the chamber. $[^{90}Y]Cl_3$ (122MBq /3.3mCi) was diluted in 0.5M $NH_4OAc$ pH 7.2 buffer and added to DOTA-hu5A10 in the same buffer, mixed gently by pipetting and incubated at 37°C for 1 hour. Radio-ITLC in 50mM DTPA was performed to confirm completion of radiolabeling. Crude $^{90}Y$-DOTA-hu5A10 was purified on a PD-10 column (GE Healthcare) and gravity eluted with 1% HSA in 0.9% saline. The radiolabeling yield was 102MBq (83% conversion) with a final specific activity of 101MBq/mg (2.73mCi/mg) and radiochemical purity of 99.6%.

***Cy5.5-hu5A10 and Cy5.5-IgG1.*** The near infrared fluorophore Cy5.5-NHS (GE Healthcare) was resuspended in methanol, aliquoted, and dried by SpeedVac. Using a 3-fold molar excess of Cy5.5-NHS, hu5A10 or an isotype control IgG1 antibody was labeled and the pH adjusted to 8.5 with $Na_2CO_3$. The reaction was shaken at 22°C for 4h followed by size exclusion gel purification on a PD10 column (GE Healthcare) and ultrafiltration (Amicon Ultra 4, Millipore). The number of dye molecules per antibody was quantified spectrophotometrically and calculated to be 1.3 (SpectraMax M5, Molecular Devices). The dye-labeled antibody conjugate was freshly prepared for each experiment.

**PET/CT**

**PET.** List-mode data were acquired using a γ-ray energy window of 350-750 keV and a coincidence timing window of 6 ns. PET image data were corrected for detector nonuniformity, dead time, random coincidences, and physical decay. For all static images, scan time was adjusted to ensure between 15 million and 25 million coincident events were recorded. Data were sorted into 3D histograms by Fourier rebinning and transverse images were reconstructed using a maximum a priori algorithm to a 256 x

256 x 95 (0.72 mm x 0.72 mm x 1.3 mm) matrix. The reconstructed spatial resolution for $^{89}Zr$ was 1.9 mm full-width half-maximum at the center of the field of view. The image data were normalized to correct for nonuniformity of response of the PET, dead-time count losses, positron branching ratio and physical decay to the time of injection, but no attenuation, scatter or partial volume averaging correction was applied. An empirically determined system calibration factor [in units of (mCi/mL) / (counts per second per voxel)] for mice was used to convert voxel count rates to activity concentrations. PET data were reconstructed using a 3D-filtered back projection maximum a priori algorithm using a ramp filter with a cutoff frequency equal to the Nyquist frequency into a 128 x 128 x 95 matrix.

**CT.** General CT acquisition parameters were 55kVp with a pitch of 1 and 240 projections in a spiral scan mode. The entire animal was scanned using a multiple-field-of-view procedure (with an approximate field of view of 4 cm x 4 cm x 4 cm per bed position), commonly requiring three bed positions per scan. Total scan time was about 10 minutes. A Shepp-Logan filter was used during the reconstruction process to produce image matrices with isotropic volumes of 221mm.

**Non-human primates**

All macaques had visual and auditory access to other macaques 24h per day, with the exception of the day of study. These animals were fed a complete lifecycle diet (LabDiet Monkey Diet 5037, PMI) twice daily, supplemented with fruits and vegetables, and offered water free choice by using automatic watering devices. Animals underwent biannual physical examinations as part of routine care. Animals were housed in standard stainless-steel caging. Daily environmental enrichment included rotating manipulanda (forage boards, mirrors, puzzle feeders, etc.), electronic auditory and visual enrichments, and novel foodstuffs. The animal holding room was maintained at 72±2 °F (21.5±1 °C), relative humidity between 30-70%, and a 12:12 hour light:dark photoperiod. Animals were fasted before procedures requiring anesthesia (ketamine hydrochloride (5±5 mg/kg) +/- dexmedetomidine (0.01±0.02 mg/kg), intra-muscular); drinking water was offered *ad libitum*. At study endpoint monkeys were anesthetized with ketamine hydrochloride + dexmedetomidine injection and then administered an intravenous injectable commercial euthanasia agent (390 mg pentobarbital (Euthasol, Virbac Animal Health). Death was confirmed by a sustained loss of heartbeat, a loss of corneal reflexes, and pupillary dilation.

**Pharmacokinetics of [$^{89}$Zr]hu5A10 in NHP - longitudinal PET/CT imaging**

Three healthy male Mauritian cynomologus macaques (*Macaca fascicularis*, PrimGen) of similar age (two 14 and one 15 years-old) and weight (10.7, 12.9, and 13.5kg, respectively) were selected for this study. Older adult males were preferentially sought to more accurately model sedentary middle-aged humans. The weight of males in the wild plateaus around 8kg at 12–13 years old (5). The pharmacokinetics and distribution of [$^{89}$Zr]hu5A10 was evaluated using a longitudinal PET/CT study over two weeks at the Weill Cornell Medicine – Citigroup Biomedical Imaging Center. Each animal was imaged dynamically for 60 minutes after injection, and subsequently at three more timepoints in the following ranges: 48–72h, 120–144h, and 216–312h. Due to ethical considerations, an individual could only be fully sedated and imaged once in a contiguous 24-hour time period. Animals were nil per os for 6h prior to imaging. One hour prior to imaging, the animals were sedated, intubated and stabilized under 2% isoflurane/oxygen anesthesia. They were then catheterized in the saphenous vein with a heparinized intravenous port for radiotracer injection. The NHP was then brought to the imaging suite, supplied with intravenous saline (brachial vein), wrapped in a blanket with heating pads. A Biograph64 mCT PET/CT (Siemens Medical Solutions) was used to image the animals positioned transversely in the bore such that the head, torso, abdomen and pelvic area were all in the PET field of view (78cm x 22.1cm). A scout CT was used to ensure proper positioning, then an attenuation correction CT was obtained. The CT acquisition was performed in helical mode using 12kV, 10 mAs and 45 x 512 x 512 matrix with a 78cm bore diameter. Each animal received a single intravenous bolus injection containing 0.20±0.02mg/kg antibody dose and 115±11MBq (3.1±0.3mCi) activity of [$^{89}$Zr]hu5A10. Saline (3mL) was used to flush the dose syringe and lines. PET data acquisition was started simultaneously with radiotracer injection and collected for 90 minutes. Animals were maintained under 2% isoflurane/oxygen anesthesia during the scanning and continually monitored for body temperature, heart rate and respiration. After scanning, the animals were extubated immediately, returned to housing, fed and monitored for several hours during recovery. For follow-up scans on subsequent days, the animals were repositioned in the same orientation and imaged for 30-60 minutes. Listmode data was sorted into histograms by Fourier re-binning and the binned data reconstructed using the ordered-subset expectation maximization (OSEM) algorithm. Reconstruction was performed iteratively with attenuation, scatter, and other standard corrections applied. The 60-minute dynamic study data were binned as follows: 6x5 sec, 3x10 sec, 3x20 sec, 2x30 sec, 4x1 min, 4x2 min, 5x5 min, 3x10 min. Using co-registered PET/CT images, volumes of interest (VOIs) were drawn around the entirety of smaller organs (spleen, L1-L4 vertebrae, kidney, prostate, and epididymis) or a spherical VOI drawn in the center of larger organs (liver, heart, intestine, thigh muscle, and testicle) with Hermes Hybrid Viewer (Hermes GoldLx v2.3.0). VOIs were not drawn in organs showing negligible uptake, such as brain, lung and salivary glands. To estimate initial blood input function, a VOI was drawn around a 4cm cylindrical segment of the femoral vein using early (0-30 seconds) PET frames from the dynamic study. The average decay-corrected activities in each VOI were converted to standardized uptake values (SUV) by means of the following calculation: SUV = activity concentration (kBq/mL)/ (injection dose (kBq)/body weight (g)). Time-activity curves (TAC) for dynamic analysis were generated using the uptake values for each VOI per frame; later timepoints were appended as single data points. SUV values were then plotted as a function of time to describe the kinetics of distribution and clearance.
