## Supplementary Table S1 for "PSA-targeted Alpha-, Beta- and Positron Emitting Immuno-Theranostics in Murine Prostate Cancer Models and Non-Human Primates"

**Supplementary Table S1. Dosimetry of [$^{225}$Ac]hu5A10 (11.1 kBq/500nCi) and [$^{90}$Y]hu5A10 (18.5 MBq/500µCi) in the LNCaP-AR s.c. model.**

| | [$^{225}$Ac]hu5A10 | | [$^{90}$Y]hu5A10 | |
|---|---|---|---|---|
| **Organs** | **Absorbed dose (Gy)** | **Absorbed Dose/IA (Gy/MBq)** | **Absorbed dose (Gy)** | **Absorbed Dose/IA (Gy/MBq)** |
| **Intestine** | 0.2 | 18.02 | 1.2 | 0.22 |
| **Spleen** | 2.77 | 249.55 | 3.26 | 0.59 |
| **Liver** | 1.98 | 178.38 | 5.36 | 0.97 |
| **Kidney** | 0.44 | 39.64 | 2.19 | 0.40 |
| **Lungs** | 0.77 | 69.37 | 2.98 | 0.54 |
| **Heart** | 0.38 | 34.23 | 2.61 | 0.47 |
| **Tumor** | 9.3 | 837.84 | 13.37 | 2.41 |

IA – injected activity
