## Supplementary Figure S1 for "PSA-targeted Alpha-, Beta- and Positron Emitting Immuno-Theranostics in Murine Prostate Cancer Models and Non-Human Primates"

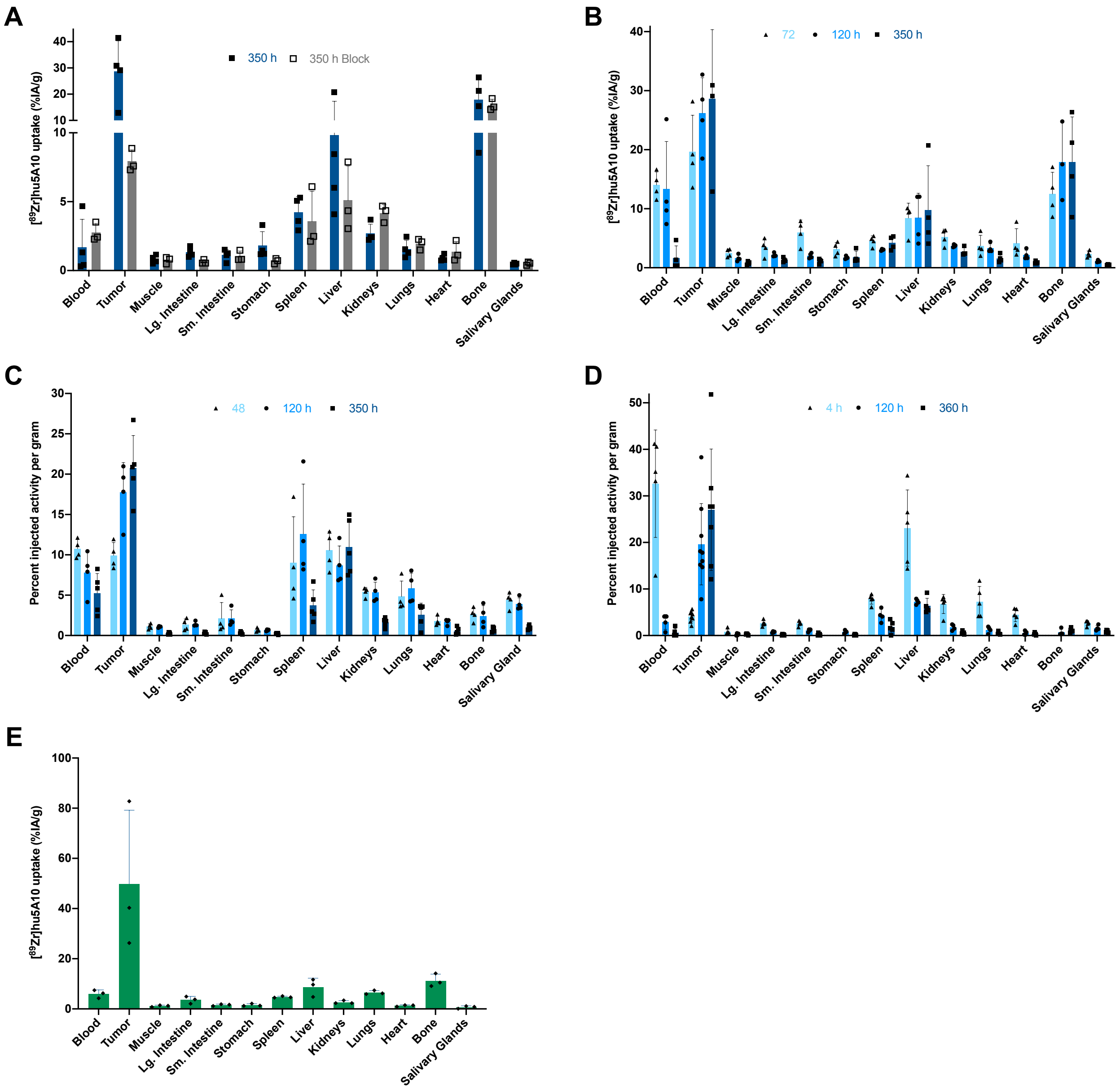


**Supplementary Figure S1. *Ex-vivo* biodistribution of [$^{89}$Zr]hu5A10.** (**A**) Specificity of hu5A10. Mice with LNCaP-AR tumors were injected with 4.07MBq/110µCi [$^{89}$Zr]hu5A10 without or with 1 mg cold antibody (block; n=3-4). (**B-D**) Biodistribution of [$^{89}$Zr]hu5A10 (4.07MBq; 110µCi; **B**), [$^{90}$Y]hu5A10 (5.55MBq; 150µCi; **C**), and [$^{225}$Ac]hu5A10 (11 kBq; 300nCi; **D**) over time in the LNCaP-AR tumor (n=3-5 per time point, except for [$^{225}$Ac]hu5A10 tumor uptake: n=7-9 tumors). (**E**) Biodistribution in MDAPCa2b xenografts at 120h p.i. of 4.07MBq/110µCi [$^{89}$Zr]hu5A10. Columns represent mean and error bars SD. Sm. Intestine – small intestine, VP – ventral prostate
