## Supplementary Figure S2 for "PSA-targeted Alpha-, Beta- and Positron Emitting Immuno-Theranostics in Murine Prostate Cancer Models and Non-Human Primates"

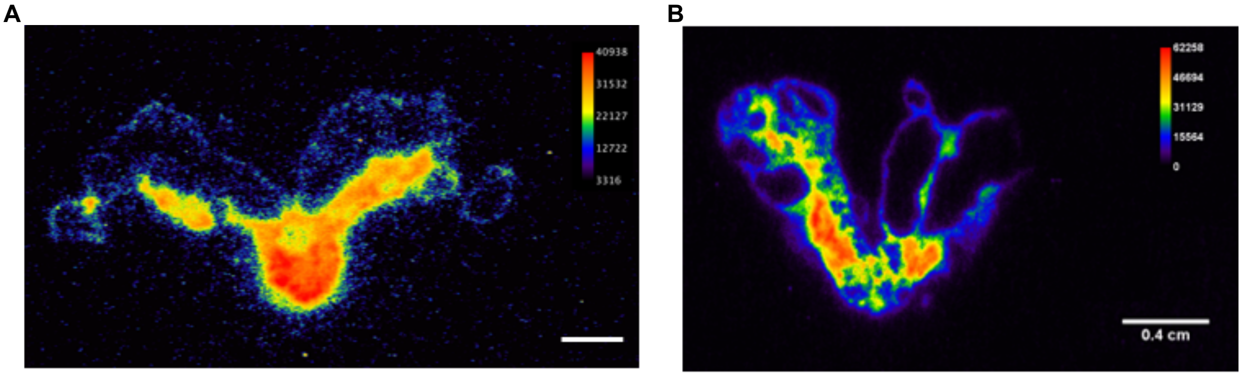


**Supplementary Figure S2.** Autoradiography of whole mounted *KLK3*_Hi-*MYC* GEMM prostate sections obtained 120h p.i. of [$^{89}$Zr]hu5A10 (**A**) or [$^{90}$Y]hu5A10 (**B**). Radioimmunoconjugate activity was detected in regions expressing PSA, but not in adjacent tissues lacking PSA expression, such as seminal vesicles and urethra.
