## Supplementary Figure S3 for "PSA-targeted Alpha-, Beta- and Positron Emitting Immuno-Theranostics in Murine Prostate Cancer Models and Non-Human Primates"

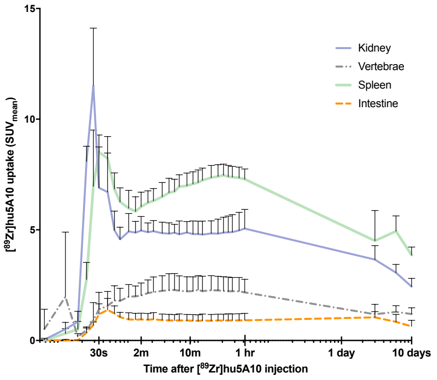


**Supplementary Figure S3.** Male cynomolgus macaque were intravenously administered 115±1MBq (3.1±0.3mCi) of [$^{89}$Zr]hu5A10 at 0.2 mg/kg for longitudinal immune-PET imaging. Time-activity curves of [$^{89}$Zr]hu5A10 for kidney, spleen, bone (vertebra) and intestine (n=3). SUV$_{mean}$ (mean+range) were calculated for each organ VOI and plotted over time.
